# Illusory path configurations reveal age-related differences in egocentric pointing variability

**DOI:** 10.64898/2026.05.06.722714

**Authors:** Abhilasha Vishwanath, Mathew F. Watson, Melanie K. Gin, Yu Du, Robert C. Wilson, Arne Ekstrom

## Abstract

A consistent finding across studies with older adults is that they typically perform worse at spatial memory tasks, particularly those conducted in virtual reality and involving novel environments, compared to young adults. While the underlying reasons for this difference remain unclear, some proposed hypotheses include differences in sensory cue integration and cue conflict resolution. Here, we tested older (n = 29) and young adults (n = 28) in immersive and walkable virtual reality using both correctly rendered and illusory hallways to test how visual cues (i.e., an intersection) and self-motion cues are integrated. In the illusory or false-intersection condition, we hypothesized that participants who walked an uncrossed path would merge two disconnected intersections, creating the illusion of a crossed path. The overall accuracy and pointing patterns were similar between young and older adults in both true- and false-intersection conditions. We did find, however, a significant age by condition interaction effect in egocentric pointing variability where older adults showed lower variability in the illusory condition and higher variability in the control condition. At the same time, older adults also drew worse maps for the control condition compared to young adults. However, the pointing error correlated with the accuracy of maps drawn regardless of age, suggesting that the pointing patterns shown by both age groups related to their underlying representations of the paths. Our findings are inconsistent with a global deficit in allocentric navigation or path integration and instead suggest that more subtle differences in strategy use might manifest with age.

## Introduction

Knowing where you are when navigating involves integrating the information provided by self-motion (proprioception, motion perception, and visual flow) and visual landmark cues (external and stable features in the environment). For example, when walking in a new building, you may be able to track your position along multiple hallways, although looking at distinctive landmarks, like intersections, may help update or reorient where you think you are (Waller & Hodgson, 2006). The influence of self-motion and landmark cues on spatial orientation and position tracking is complex, particularly given noisy self-motion information and during uncertain landmark cues (Etienne et al., 2004; Harootonian et al., 2020; Sjolund et al., 2018; Stangl et al., 2020). Some past studies have looked at situations in which self-motion and landmark cues conflict, which have typically reported that landmarks can override less reliable self-motion cues, although these have typically been for simple trajectories such as straight-line trajectories and triangle completion tasks (Chen et al., 2017; Nardini et al., 2008; Petzschner & Glasauer, 2011; Zhao & Warren, 2015). Here, we wished to better understand the situations in which landmarks, such as intersections in Hallways with varying geometries, compete with self-motion cues and what, if any, changes occur with age.

Older adults typically perform worse at spatial memory and navigation tasks than young adults. Some studies have suggested that older adults may perform worse at path integration, which involves combining self-motion cues during translations and rotations to estimate one’s current position, that could in turn contribute to their worse performance on navigation (Adamo et al., 2012; Allen et al., 2004; Stangl et al., 2018, 2020). Age-related differences in path integration could stem from increased neural noise in areas such as the entorhinal cortex (Lester et al., 2017; Segen et al., 2022; Stangl et al., 2018), and greater noise accumulation in velocity estimates (Stangl et al., 2020). Another possible explanation for increased errors in navigation could be alterations in how self-motion and visual cues are combined in older adults (Allen et al., 2004; Campbell et al., 2018; Hill et al., 2025; Shayman et al., 2024). Specifically, older adults might be weighting self-motion and visual landmark cues differently, or possible sub-optimally, compared to young adults (Bates & Wolbers, 2014; Shayman et al., 2024, 2025). For instance older adults, compared to younger adults, over-weight uncertain visual cues more than stable visual cues (Shayman et al., 2025). By testing older and young adults under situations of competing path integration and landmark information, we can better determine the extent to which age-related navigation differences might stem from worse path integration performance or worse combination of path integration and landmark cues.

Another possible explanation for differences in spatial navigation performance in older adults is that they have a selective impairment in allocentric navigation with relatively preserved egocentric navigation in novel environments. This explanation comes from findings that older adults sometimes show difficulty in finding targets from new locations/orientations but have little difficulty in finding these targets from the learned start locations/orientations (Fernandez-Baizan et al., 2020; Merhav & Wolbers, 2019; Moffat & Resnick, 2002). In contrast, some studies have reported no such difference in older adults for repeated location (putative egocentric navigation) compared to a new location (putative allocentric), but rather a worse overall performance in older adults (Hill et al., 2024; McAvan et al., 2021). Thus, such “deficits” in allocentric navigation may instead stem from a weak preference for egocentric strategies such that older adults may employ familiar and learned routes more often than young adults, possibly due to rigidity in strategy use (Hill et al., 2025). Other studies have highlighted this decreased flexibility, in which older adults show worse performance compared to young adults when switching between putative egocentric and allocentric strategies, also consistent with potential inflexibility or rigidity in behavior (Harris et al., 2012; Lemay et al., 2004; Yamamoto & DeGirolamo, 2012).

To answer these age-related questions about conflicting sensory cue conditions and rigidity in older adults we developed an immersive virtual reality task with illusory intersections along walked paths. The task is adopted from a previous work by Du et al., (2023), in which the traversed paths consisted of ‘Hallways’, in immersive and walkable virtual reality, involving visual illusions in the form of false ‘+’ intersections. For example, the illusory paths would not cross over in physical space but the participants were shown false ‘+’ intersections to induce an illusion of a crossing path while the control paths would involve seeing the ‘+’ intersection in the same location, as would happen during real-world navigation. Thus, the false or illusory intersection created a mismatch between the self-motion and visual landmark cues, in which the walked paths differed from the visual intersections that could not exist in Euclidean space.

In the current study, we modified this Hallways design to have better visibility of the intersections and better juxtapose the self-motion and visual landmark cues. A cohort of both young and older adults were tested in the two conditions, involving uncrossed paths with an illusory intersection and control paths in which the intersection was rendered correctly, with accompanying map-drawings of traversed paths in the two conditions. Based on the previous work suggesting that an illusory intersection can produce distortions in spatial memory, we hypothesized that the illusory intersection might elicit a ‘Cross Illusion’ in terms of pointing back to the start location. We also predicted age differences in the patterns of pointing, although these differed based on the hypotheses mentioned above. One prediction, based on the hypothesis of a global “deficit” in path integration with age, is that older adults would rely more on the landmark cues, i.e., intersection, given greater noise or uncertainty in self-motion information and thus show greater cross illusion like pointing error. Alternatively, based on a difference in sensory cue integration, older adults might adopt sub-optimal approaches when resolving real and illusory intersections (Bates & Wolbers, 2014; Shayman et al., 2024) and resort to averaging or regression to the mean egocentric pointing. Another possible prediction, based on the allocentric deficit with age hypothesis outlined above, is a decreased overall accuracy in the map-drawings and reduced correlation between the egocentric pointing and rendered drawings in older adults.

## Methods

### Participants

30 younger adults (14 males, mean age 22.63 years) and 31 older adults (12 males, mean age 71.16 years) were recruited for the study from the University of Arizona and surrounding Tucson city area in the year 2025. Participants were compensated with either $20/hour or course credit. The Institutional Review Board at the University of Arizona approved the study and methods. All participants provided written informed consent before participating in the study.

The breakdown of the race/ethnicity of the participants is as follows, young adults: African American (n = 2), Asian (n = 7), White (n = 12), Hispanic (n = 2), two or more race/ethnicities (n = 5), un-declared (n = 2); older adults: Asian (n = 1), White (n = 27), Hispanic (n = 2), un-declared (n = 1).

The Older adults were screened for cognitive decline characteristic of the early stages of dementia using the Jak/Bondi criteria (Bondi et al., 2014). Briefly, a battery of neuropsychological tests assaying verbal memory, visual memory, executive function, language fluency, and other neuropsychology constructs was employed. Any older adult performing 2 standard deviations below the mean on any (or more) two tests within a domain or 3 (or more) tests across domains was considered to have mild cognitive impairment. These individuals were not included in the study due to concerns about potential early dementia.

One participant (older adult, female) was excluded from the study due to inability to complete the experiment. Three additional participants (2 young and 1 older adult) were classified as outliers (using Grubbs’ test) in pointing behavior for the control condition and were excluded from the analysis. This sample size of 28 young and 29 older adults, based on the study by Du et al., (2023) is powered to detect moderate-to-large within-subject effects (*d* > 0.6) between control and illusory conditions with approximately 80% power.

### Apparatus

The Hallways task was fully developed using Landmarks 2.0 framework (Starrett et al., 2021) in Unity 2018.4.11f1 as previously described in Du et al., (2023). The virtual environment spanned approximately 5 m by 5 m and was rendered using wireless HTC Vive pro (HTC, New Taipei City, Taiwan) headset and controllers (Fig. 1A left).

**Figure 1:**
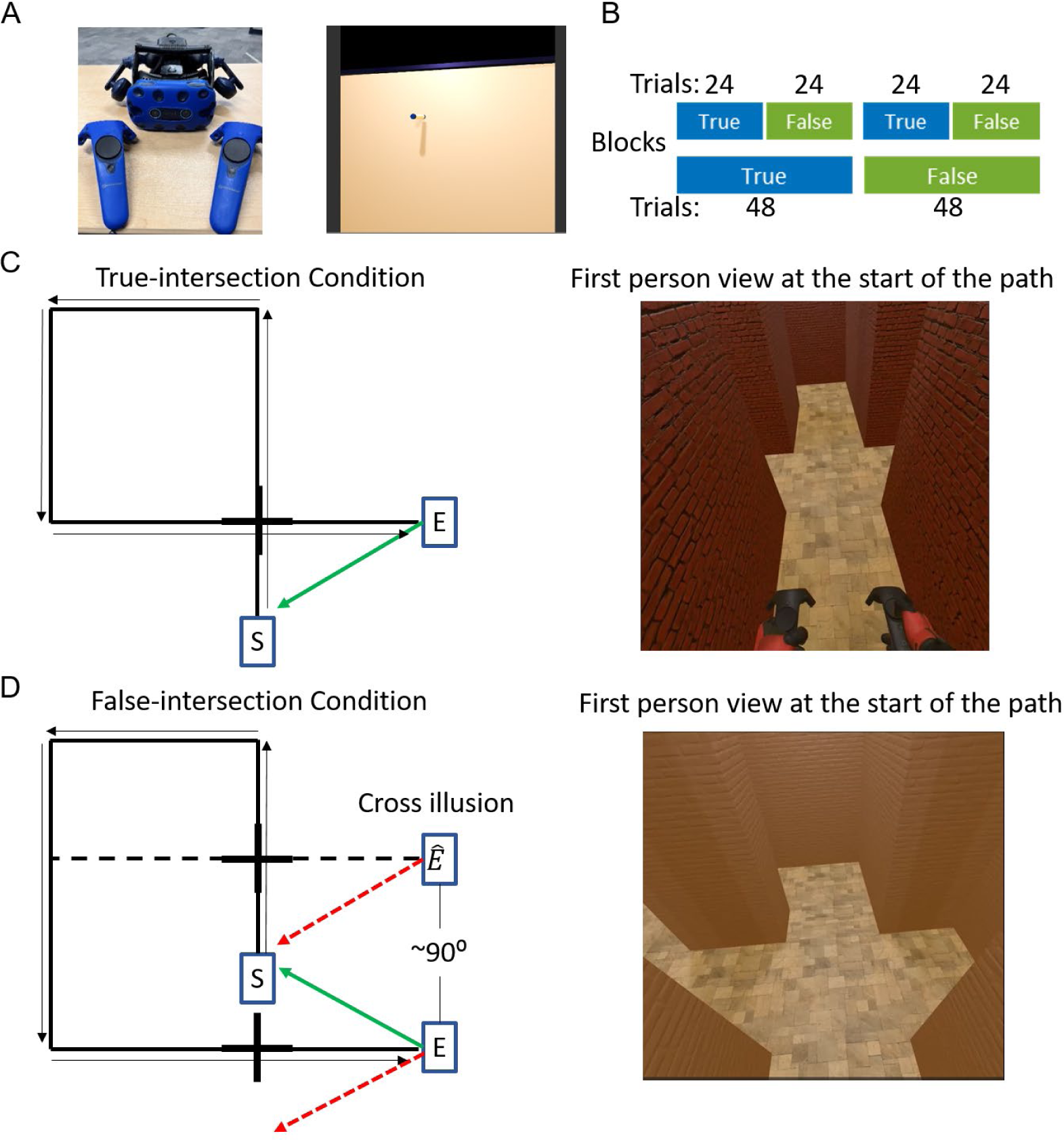
Experimental design and Hallway task conditions. **A)** The HTC VIVE handheld controllers and headset used to perform immersive virtual reality task (LEFT). First person view of the virtual whiteboard and virtual marker that was used by participants to draw maps of the walked paths (RIGHT). **B)** Participants completed both true-intersection and false-intersection conditions in 48×2 or 24×4 trial blocks, which were counterbalanced across participants, for a total of 96 trials. **C)** Top-down schematic of a representative path configuration for the true-intersection condition (LEFT), with start and end locations marked at ‘S’ and ‘E’ respectively. The solid green arrow shows the correct pointing response back to the start location. The black arrows show the continuous trajectory that participants take on each trial. RIGHT: the first person view in virtual reality at the start of a true-intersection condition path. **D)** Same as in C) but for false-intersection condition. The dashed red lines in the top-down schematic shows the predicted Cross Illusion pointing pattern to the start location where the dashed black line shows the illusion of a crossed path due to the false-intersections.

### Design

The hallway layouts were designed to have either a true intersection or a false intersection, which were created by either accurately rendering a visible intersection or showing a physically impossible non-intersecting intersection. The top-down and first-person view of these types of hallways are shown in Figure 1C and D. In every hallway path, participants encountered intersections twice, at the start and towards the end of their traversed path. In the true-intersection condition, participants crossed over their path and saw the real intersection at two instances, which reflected their walked trajectory in physical space (Fig. 1C left).

In the false-intersection condition, participants were shown an illusory intersection at two locations as if making an actual cross intersection, but unlike the control condition, these paths did not intersect in physical space (Fig. 1D left). The overall lengths of these paths were matched across true- and false-intersection conditions. To generate the illusion of a false intersection in the illusory condition, the hallway paths at the first intersection resembled true-intersection paths (Fig. 1C and D right). But, the hallway configuration changed in real time at the second turn in these false-intersection hallway paths, outside participant’s view, to make a seamless visual depiction of a crossed path at the second intersection encounter. Thus, in false-intersection paths the path integration estimates did not match the visual intersections, which served as landmarks in these paths. We hypothesized that the false-intersection condition would induce a cross illusion, i.e., ∼90 degree angular pointing error (Fig. 1D left).

### Procedure

There were 8 unique paths in the two conditions (true-intersection and false-intersection) which comprised of mirrored left and right turning paths (n = 4 each). Each set of left and right paths were generated with different lengths of the 4 legs of the paths, keeping the total length walked consistent for each path. All participants experienced both conditions in a blocked design (Fig. 1B) counterbalanced across participants. Thirty-nine out of 57 participants experienced all true-intersection trials followed by all false-intersection trials (Fig. 1B). The remaining participants experienced alternating true- and false-intersection trials in 24 trial blocks. Within each block, the 8 unique paths were pseudo-randomized for each participant. Ninety-six trials were completed per participant (48 trials per condition), and each unique path was encountered a total of 6 times. Seven participants out of the 57 total participants completed a total of 48 trials (24 per condition), with each unique path encountered 3 times.

Participants began each trial with a distractor task during which they rotated in darkness while counting backwards from a set of numbers. An example instruction message read, “Count backwards from 105 by 3”. Next, participants were guided, in the dark, to a starting location indicated by a marker on the virtual floor. The starting location was randomized between four different locations. Once at the starting location, the virtual environment with the hallways were shown (Fig. 1C and D right) and participants were asked to follow the hallway to the end. They were instructed to look through the intersections as they moved through the hallways. A researcher was present at all times to make sure participants kept to the path indicated in Fig. 1 C and D. Once at the end of a hallway/path, all visual input was removed and participants were asked to use their controller and point to their starting location as accurately as possible. A virtual laser beam extending from one of the hand-held controllers was used to mark the starting location. Participants were asked to take into account both the distance and direction when indicating the start location. Both direction/angular and distance error from the target were logged and subsequently analyzed.

### Map drawing and Procrustes distance analysis

On ∼25% of trials, participants were asked to draw the path walked on the corresponding trial using a virtual whiteboard (Fig. 1A). Participants who did not complete at least 60% of map-drawing trials were excluded from analysis. One older adult was excluded this way from the map-drawing analysis.

The hand drawn maps of the paths were converted into coordinates using MATLAB such that the x and y coordinates of the start, end, and all turns were logged. The dissimilarity between map drawings and the actual path walked for a given trial was given by the Procrustes distance analysis. This analysis performs an optimal transformation between the two shapes and provides the distance, i.e., dissimilarity, following the transformation and the actual path. This was implemented using the ‘procrustes’ function in MATLAB.

### Data analysis

Angular pointing error was calculated as deviation between participant’s response and the correct pointing angle from the end position in a given hallway/path. Positive and negative angular pointing error represented deviation to the right and left side of the correct angle respectively. Distance error was calculated as the Euclidean distance between the start location and the location of the participant’s response. The mean of signed pointing error (or angular pointing error) and standard deviation of the pointing error (or variable pointing error) were obtained using circular statistics (MATLAB, Circular statistics toolbox, circ_mean and circ_var functions). The pointing, distance and standard deviation of pointing errors were averaged across the left and right turn hallway paths and all trials for a given unique path for each participant unless otherwise reported.

There was no significant difference in angular pointing error in the false-intersection condition between 24×4 and 48×2 block types (Two-sample t-test, *p* = 0.262) or between participants who completed 24 and 48 total trials (Two-sample t-test, *p* = 0.401). Thus, the analysis reported in the results section is averaged across participants regardless of block type and total number of trials.

K means clustering (using kmeans function MATLAB) was used to divide the pointing responses, converted into x and y coordinates, into three subgroups. A mixed effects linear regression model was applied to analyze the condition (true-intersection and false-intersection) and age (Young and Old) effects with the participants as random effects accounting for the within subject design. Subgroups in false-intersection condition were determined using a k-means clustering approach. A 2×3 factorial ANOVA was used to compare the false-intersection condition subgroups (Accurate, Averaging and Cross Illusion) and age effects.

## Results

Our primary goal was to determine whether conflicting visual and path integration cues altered the mental representation of the physical geometry in the paths walked by participants. A secondary goal involved testing whether this would vary by age and if so, how. The pointing response for all participants in an example path in true-intersection and false-intersection conditions are shown in Figure 2A-B for both young and older adults. The pointing response for the other unique paths in the two conditions are shown in supplementary figure 1.

**Figure 2:**
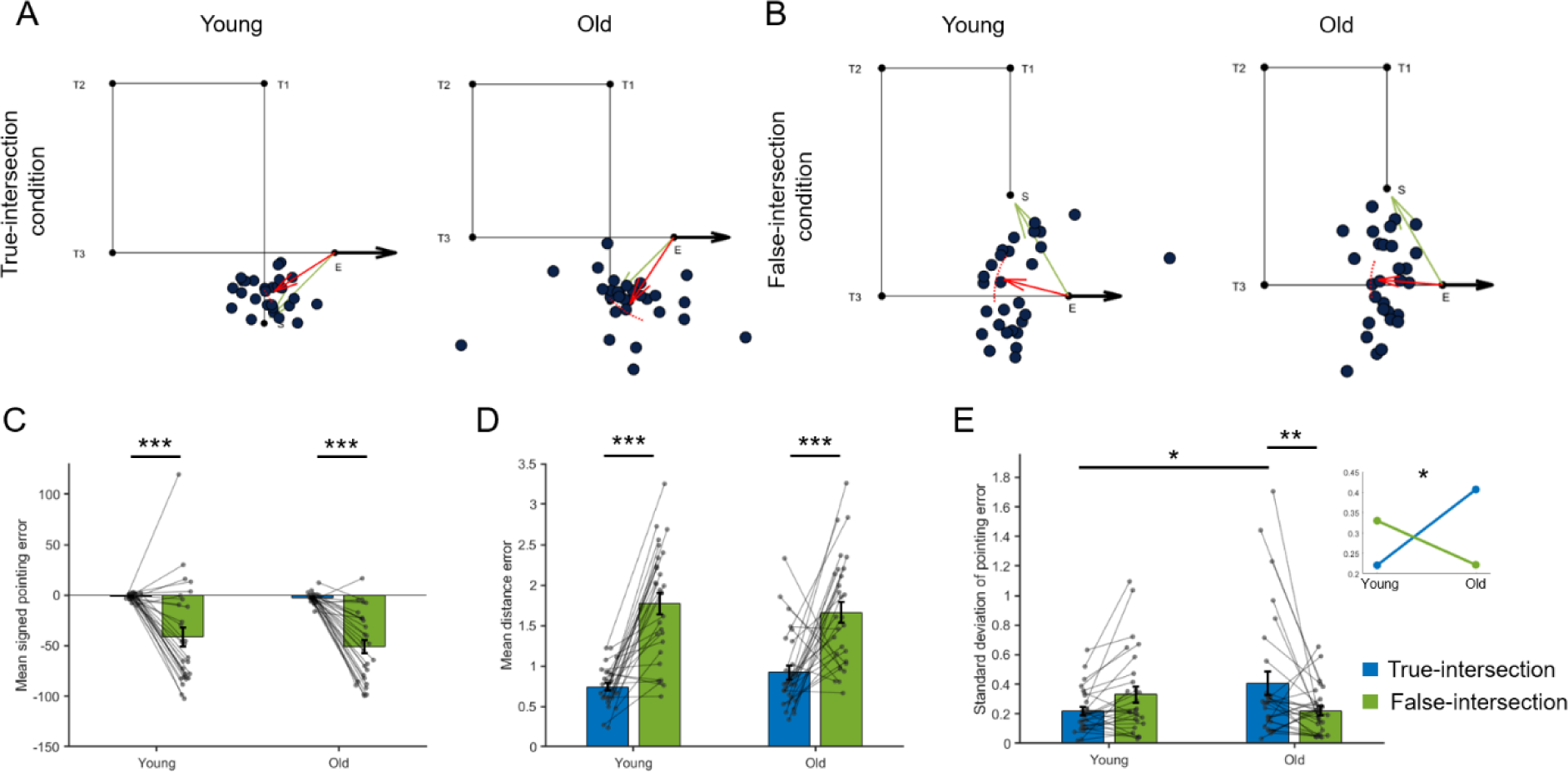
Pointing response in the two Hallway conditions. **A)** Participant’s response location (blue dots) on the true-intersection condition averaged across trials for a representative path configuration in young (left, n = 28) and older adults (right, n = 29). The ‘S’ and ‘E’ represent the start and end points for the path. The ‘T1’, ‘T2’, and ‘T3’ represent the three turns for the given hallway path. **B)** Same as in A), but for the false-intersection condition. **C)** Average signed pointing error across participants compared between true- and false-intersection conditions for young (n = 28) and older adults (n = 29). **D)** Same as C), but for distance error. **E)** Same as C), but for the standard deviation of pointing error. INSET: Mean standard deviation of pointing error for young and older adults in true- and false-intersection conditions, showing a significant interaction effect. Error bars show ±SEM. * *p* < 0.05, ** *p* < 0.01, *** *p* < 0.001 from a linear mixed effects model.

### False intersections elicit greater angular and distance errors across age groups

The true-intersection and false-intersection conditions produced significantly different patterns of pointing in both angular and distance estimates (Fig. 2C-D). A linear mixed effects regression model was conducted to examine the effects of condition (true-intersection and false-intersection) and age (Young and Old), and their interaction on angular pointing and distance errors. The model included random intercepts and random slopes for condition within subjects. There was a significant main effect of condition on angular pointing error (Fig. 2C; *b* = 0.84, SE = 0.14, *t*(110) = 6.07, *p* < 0.001), indicating higher signed pointing error in the false-intersection condition compared to true-intersection condition. However, there was no main effect of age group (*b* = 0.16, SE = 0.19, *t*(110) = 0.83, *p* = 0.408) or interaction (*b* = -0.14, SE = 0.2, *t*(110) =-0.71, *p* = 0.48). Similarly, distance error also showed a main effect of condition (Fig. 2D; *b* = - 0.74, SE = 0.15, *t*(110) = -4.82, *p* < 0.001) and no main effect for age group (*b* = 0.11, SE = 0.18, *t*(110) = 0.6, *p* = 0.547) or interaction (*b* = -0.29, SE = 0.22, *t*(110) = -1.3, *p* = 0.195). Therefore, the false-intersection condition significantly altered angular and distance response of participants regardless of age.

### Variability in pointing error differed across conditions but only in older adults

We compared variable pointing error across age groups and conditions to assess whether the conflicting visual and path integration cues a produced greater spread of pointing angles. A linear mixed effects regression model for variability (given by standard deviation) of pointing error showed a significant main effect of condition (Fig. 2E; *b* = 0.18, SE = 0.06, *t*(110) = 2.97, *p* = 0.003). The model also revealed a significant interaction of condition and age (*b* = -0.29, SE = 0.08, *t*(110) = -3.32, *p* = 0.001), indicating that the effect of condition on standard deviation of pointing error was different across age groups. However, there was no main effect of age group (*b* = 0.11, SE = 0.06, *t*(110) = 1.82, *p* = 0.072). Follow-up contrasts for true-intersection condition between age groups showed significantly higher variable pointing error for older adults compared to young adults (*p* = 0.029), while there was no significant difference between age groups for the false-intersection condition (*p* = 0.072). Overall, older adults had greater variability in the stable true-intersection condition and significantly lower variability in the conflicting false-intersection condition. In contrast, young adults showed an opposite trend between the two conditions. These findings suggest age differences in egocentric pointing variability and some subtle strategy rigidity in older adults. We probe the ideas of both greater variability in egocentric updating and strategy rigidity related to age in the next sections.

### False intersections elicited distinct groups of pointing patterns

We further investigated the false-intersection condition to understand the patterns of pointing elicited across the age groups (Fig. 1B). Based on our predictions for the false-intersection condition, we segmented participants into three subgroups. We subjected the participant’s pointing data averaged across all false-intersection paths from both young and older adults to k-means clustering (Fig. 3A). We classified three subgroups that had significantly different angular pointing responses (Fig. 3B-C). The subgroups were labeled Accurate, Averaging, and Cross Illusion based on the pointing error for each group. The accurate subgroup had the lowest mean error (Young, n = 8, mean = 4.7 degrees; Old, n = 10, mean = -11.8 degrees) indicating mostly accurate pointing responses. The cross illusion subgroup had the highest mean error (Young, n = 11, mean = -84.3 degrees; Old, n = 10, mean = -88.7 degrees) which was close to the predicted ∼90 degree error for the cross illusion response. Lastly, the averaging subgroup showed mean error between the two extremes (Young, n = 9, mean = -29.8 degrees, Old, n = 9, mean = -51.5 degrees). Importantly, the subgroups determined this way showed comparable number of young and old participants in each subgroup. Thus, there was a similar distribution of participants for a given pointing pattern in young and older adults.

**Figure 3:**
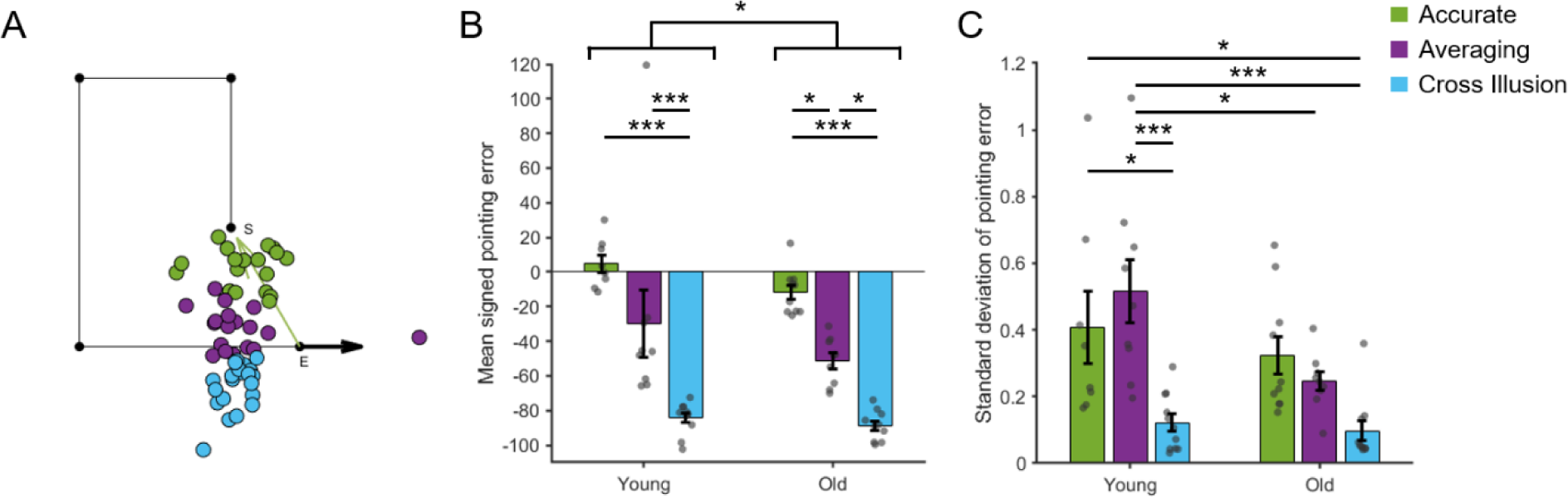
Distinct patterns of pointing emerge in the illusory intersection condition. **A)** Average participant’s response location across all paths in the false-intersection condition for both young and older adults. The colored dots show the response in the three subgroups: Accurate (green), Averaging (purple), and Cross Illusion (blue). **B)** Average signed pointing error across participants compared between the three subgroups for young and older adults. **C)** Same as B), but for the standard deviation of pointing error. Error bars show ±SEM. * *p* < 0.05, ** *p* < 0.01, *** *p* < 0.001 from a 2×3 ANOVA.

The angular/signed pointing error showed a significant main effect of subgroups (Fig. 3B; ANOVA, F(51) = 52.27, *p* < 0.001) and a significant, albeit modest, main effect of age group (F(51) = 4.42, *p* = 0.04). Post-hoc comparisons did not show significant differences between subgroups in young and old adults, and therefore the main effect of age on signed pointing error was likely driven by differences across the subgroups. This overall greater pointing error bias in older adults might suggest a weak tendency to regress to the midway point between the accurate and cross illusion angles in older adults.

As before, variable pointing error showed significant differences across the false-intersection subgroups and age. The subgroup by age group ANOVA showed a significant main effect of subgroup (Fig. 3C; ANOVA, F(51) = 13.03, *p* < 0.001) and age (F(51) = 6.35, *p* = 0.015) but no interaction (F(51) = 2.25, *p* = 0.116). There were no significant differences between subgroups in older adults; in young adults, post-hoc comparisons showed that the cross illusion subgroup had the lowest variable pointing error compared to accurate (*p* = 0.022) and averaging (*p* < 0.001) subgroups. Post-hoc comparisons between age groups also showed that the averaging subgroup in young adults had significantly higher variability compared to averaging (*p* = 0.038) and cross illusion (*p* < 0.001) subgroup in older adults. The cross illusion subgroup in older adults also showed significantly lower variability compared to the accurate subgroup in younger adults (*p* = 0.013). In summary, variable pointing error was significantly different between subgroups, with older adults in the averaging subgroup showing less variable pointing error compared to young adults.

### Learning characteristics in pointing behaviors across trials in true- and false-intersection conditions

We investigated the extent to which participants learned the two different conditions over trials by quantifying how their patterns of pointing changed over the course of the experiment (Fig. 4 and 5). The trials were divided into 6 blocks, which consisted of average pointing responses of 8 trials per block, allowing us to estimate the rate of change (slope) of pointing changes. To consider how pointing error changed with learning, we regressed the magnitude of pointing error (absolute value of pointing error) against trial blocks. There was a significant negative slope in absolute pointing error across trials, i.e., improved accuracy, in the true-intersection condition for both young (One sampled T-test, *p* = 0.017, Cohen’s *d* = -0.47) and older adults (One sampled T-test, *p* < 0.001, Cohen’s *d* = -0.72). However, the true-intersection condition learning slope did not differ between young and older adults (Two sampled T-test, *p* = 0.099). This suggested that both groups showed improvement in pointing to the start location over blocks.

**Figure 4:**
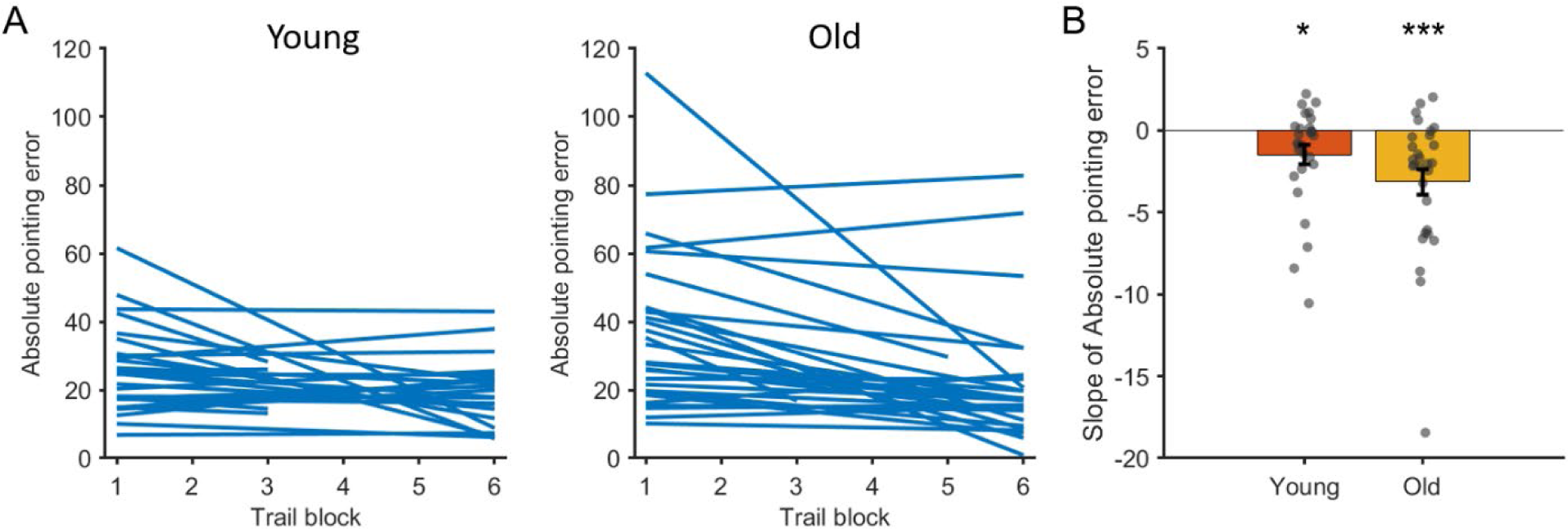
Learning slopes across trials in the true-intersection condition. **A)** Average absolute pointing error per trial block (each block comprised of 8 trials) represented as a line for each participant in young (left, n= 28) and older adults (right, n = 29). **B)** Average slope of absolute pointing error across the six trial blocks for young and older adults. Error bars show ±SEM. * *p* < 0.05, *** *p* < 0.001 from independent sample T-test.

**Figure 5:**
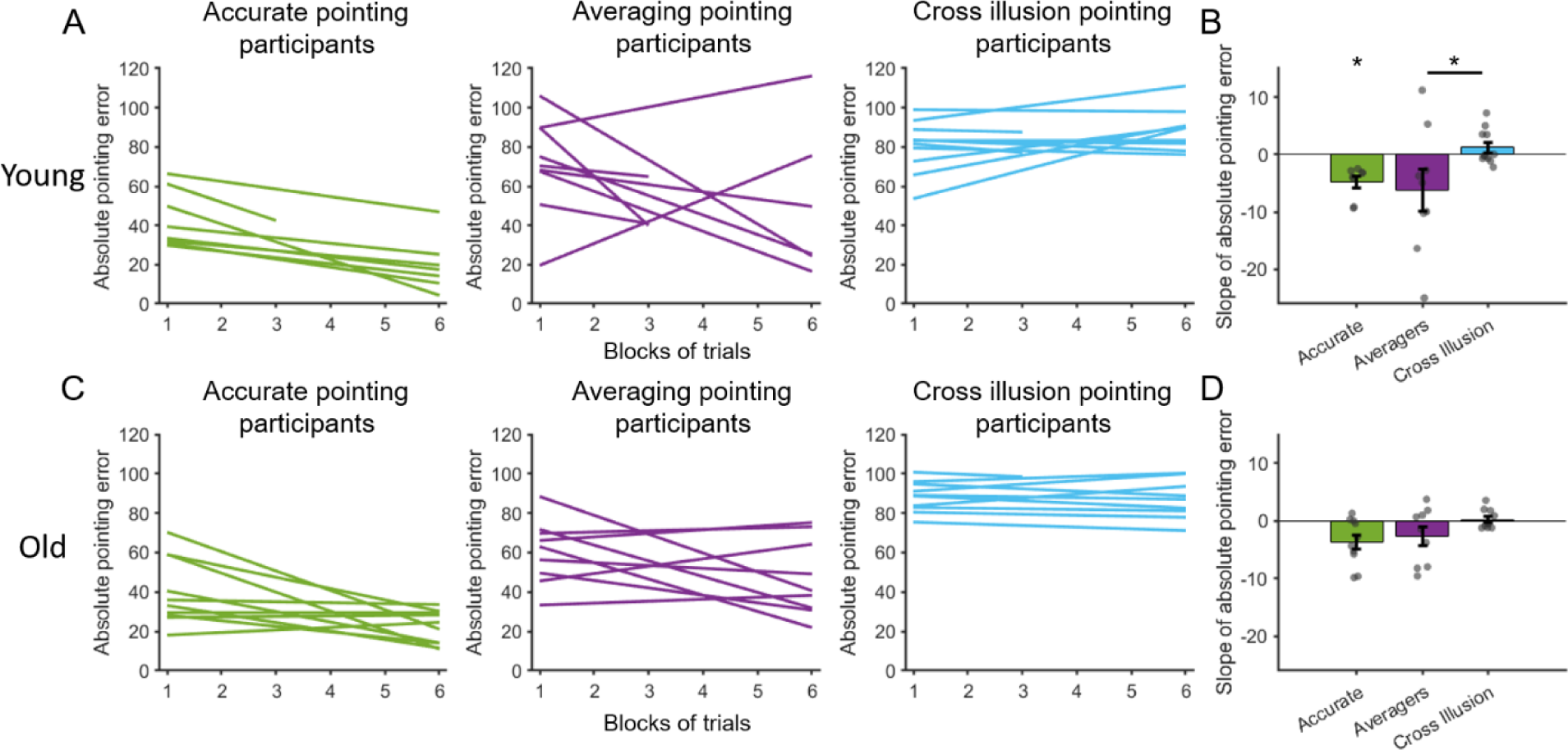
Learning slopes for the three subgroups in the illusory condition. **A)** Average absolute pointing error per trial block (each block comprised of 8 trials) represented as a line for each young adult participant in Accurate, Averaging and Cross Illusion subgroups (left to right). **B)** Average slope of absolute pointing error across the six trial blocks in adults for the three subgroups. **C-D)** Same as A-B) but for older adults. Error bars show ±SEM. * *p* < 0.05 from independent or two-sample T-test.

In the false intersection condition, the learning slope was computed for the three subgroups separately (Fig. 5). The cross illusion subgroup in both young and older adults had positive or near zero slopes (mean slope = 1.29 and -0.11 respectively). The averaging and accurate subgroups had negative slopes but only the accurate subgroup in younger adults showed a significantly negative slope (one sampled T-test, *p* = 0.012). The 3×2 ANOVA for the learning curve revealed a main effect of subgroup (F(51) = 5.67, *p* = 0.006) but no effect of age (F(51) = 0.61, *p* = 0.438) or an interaction (F(51) = 1.08, *p* = 0.348). Post-hoc comparisons showed a significant difference between the averaging and cross illusion subgroups only in the young adults (*p* = 0.03). These findings suggest that in the false intersection condition, which had conflicting cues, those susceptible to the cross illusion did not improve over trials like the true-intersection condition. Additionally, only the young adults showed any improvement in the false intersection condition, where the averaging participants improved across trials, but this was not seen in older adults. This suggests a restricted range in pointing, i.e., rigidity, even across trials during cue conflict conditions in older adults.

### Accuracy of map drawings was not distinguishable between the true-intersection and false-intersection conditions in older adults

To better understand the mental representation of the hallways and test the hypothesis that age results in a selective loss of allocentric (but not egocentric) representation, participants were asked to draw a map of the walked path on a subset of the trials. The dissimilarity between the drawings and the actual path walked was analyzed using the Procrustes distance, in which a higher distance indicates greater dissimilarity. Representative examples of participant’s drawings, the Procrustes transformation of the drawing, and the actual path walked are shown in Figure 6A. We compared the paths represented by the maps drawn with those represented by pointing to the start location to determine whether the two corresponded (Fig. 6B). There was a significant positive correlation between the absolute pointing error and the Procrustes distance in both true-intersection (Pearson’s r = 0.34, *p* = 0.0096) and false-intersection conditions (Pearson’s r = 0.52, *p* < 0.001), in which greater dissimilarity of participants’ drawings related to greater pointing error. These findings suggest that pointing error correlated with map drawing performance and that this correlation was invariant to age.

**Figure 6:**
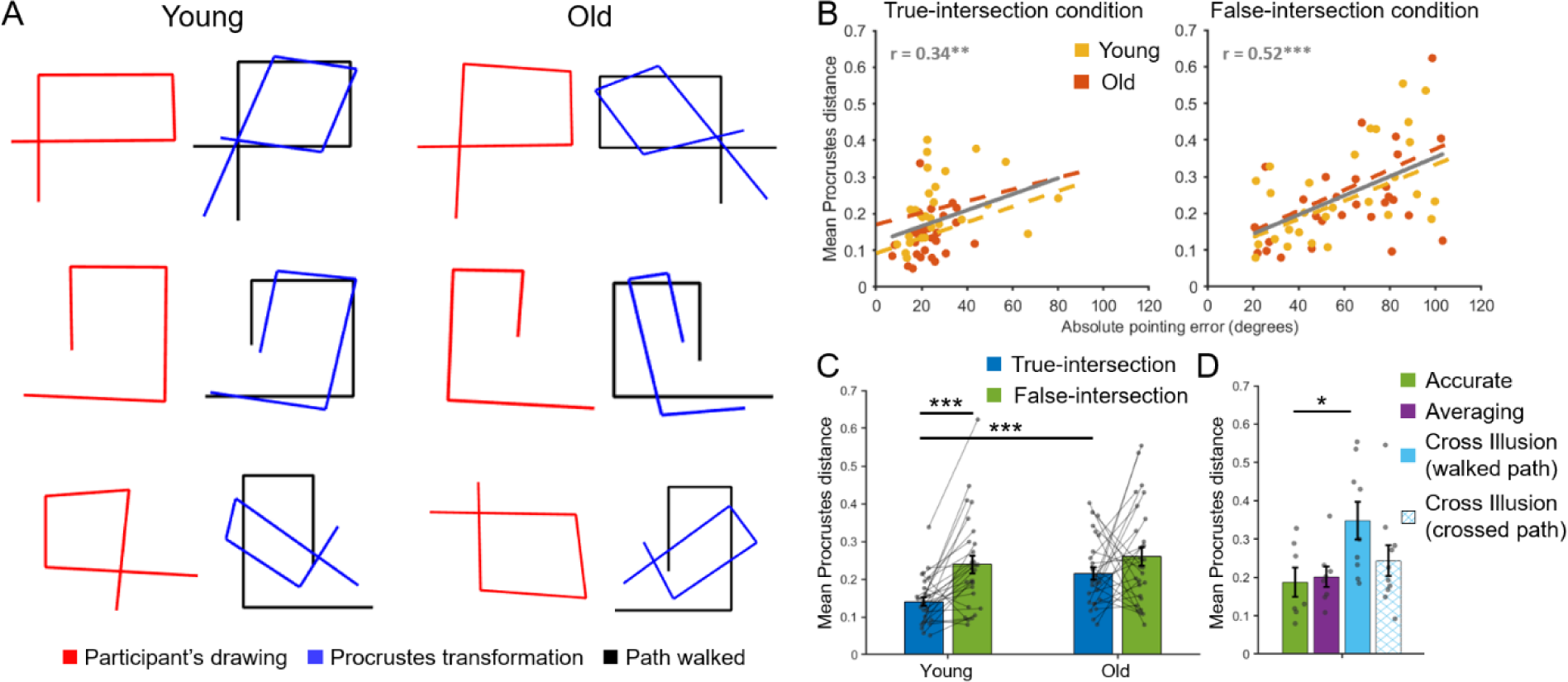
Map-drawing distortions across age groups and conditions. **A)** Representative examples of participants’ map-drawings (red), the actual path walked (black) and the Procrustes transformation of the participant’s map-drawing (blue) in young and older adults. **B)** Scatter plots showing correlation between average Procrustes distance and average absolute pointing error for each participant in true-intersection (LEFT) and false-intersection conditions (RIGHT). Each individual dot is a participant showing young (n = 28, yellow) and older adults (n = 28, orange). **C)** Average Procrustes distance across participants compared between true- and false-intersection conditions for young and older adults. **D)** Average Procrustes distance across both young and older adult participants compared across the three subgroups from the illusory condition and an additional group of cross illusion participants’ drawings compared to crossed paths instead of walked paths. Error bars show ±SEM. * *p* < 0.05, ** *p* < 0.01, *** *p* < 0.001 from Pearson’s correlation or linear mixed effects model or 2×4 ANOVA.

A linear mixed effects model comparing Procrustes distance between the conditions and age groups showed no main effect of condition (Fig. 6C; *b* = -0.04, SE = 0.03, *t*(108) = -1.69, *p* = 0.094) or age group (*b* = -0.02, SE = 0.03, *t*(108) = -0.62, *p* = 0.535) or interaction (*b* = -0.05, SE = 0.04, *t*(108) = -1.44, *p* = 0.152). Follow-up contrasts showed that the Procrustes distance for young adults in the true-intersection condition was significantly lower than the false-intersection condition (*p* < 0.001). Procrustes distance was also lower for young adults than older adults in the true-intersection condition (*p* < 0.001). There was no significant difference between true-intersection and false-intersection conditions in the older adults, nor a difference between young and older adults in the false-intersection condition. We return to greater consideration and interpretation of map drawing, particularly as a function of age, in the discussion.

### Erroneous maps drawn in false-intersection condition relate to cross-illusion representations

To better understand how those in the distorted subgroup of the false-intersection condition might represent the path they walk, we subjected the map drawings to further analysis to reveal whether the cross illusion pointing response also elicited a distorted mental representation of the walked path. We quantified the Procrustes distance for each of the three subgroups of the false-intersection condition. An additional comparison was added to this analysis, which consisted of cross illusion participants’ drawings compared to crossed paths resembling the crossed true-intersection condition, rather than the actual path walked. This comparison was included to capture whether cross illusion participants drew crossed paths, in which case the Procrustes distance in this group would be lower than the cross illusion subgroup drawings compared to actual path walked.

The 3×2 ANOVA indicated a significant effect of subgroup on the Procrustes distance (ANOVA, F(3) = 3.36, *p* = 0.024), with no age (F(1) = 0.12, *p* = 0.726) or interaction effects (F(3) = 0.41, *p* = 0.745). The mean Procrustes distance between the four subgroups, combined across age groups, are shown in Figure 6D. As predicted, post-hoc comparisons showed a significant difference between accurate and cross illusion participants (compared to actual path walked; *p* = 0.024). Importantly, though, the cross illusion participants drawings, when compared to crossed paths, were not significantly different from the other subgroups (all *p* > 0.05). Thus, these results suggests that susceptibility to the cross illusion in false-intersection condition also elicited a distorted mental representation where participants drew a crossed path (representative examples in Fig. 6A bottom row)

## Discussion

We tested spatial memory in a Hallways task using immersive and walkable VR, half of which were rendered as they would be in the physical world (true-intersection condition) and the other half of which involved an illusory intersection (false-intersection condition), to better understand cue conflict conditions in older and young adults. Our experiment was designed to test several rival hypotheses regarding spatial memory differences in older compared to young adults, including the idea that older adults have a path integration deficit, show a more rigid navigation behavior, and/or have a selective allocentric but not egocentric deficit. Our findings did not unambiguously support the predictions of any one hypothesis and can be taken to suggest that older adults show no global or obvious “deficit” in how they navigated the hallways in our task. Instead, our findings suggest that both young and older adults were susceptible to the illusion of a crossed hallway based on illusory intersection in the false-intersection condition, although different participants showed different degrees of susceptibility to the illusion.

The illusory intersection condition elicited as predicted a Cross Illusion (∼90 degree offset) but only in a subset of participants with additional subgroups showing accurate and averaging performers. These results point to the varying cue integration strategies during uncertain or conflicting landmark cues (Chen et al., 2025; Harootonian et al., 2022; Kessler et al., 2024; Zhao & Warren, 2015) that could have led to the individual differences we observed in our experiment independent of age. We can infer strategy differences in sensory cue utilization for the different subgroups, where cross illusion subgroup perceived or relied on the false intersection as a real intersection that created a crossed path. On the other hand, the accurate subgroup would need to ignore the illusory intersection and rely more on their self-motion cues to estimate their starting location. Consequently, when we looked at the map-drawings, the drawings from the cross illusion subgroups resembled a crossed path while the accurate subgroup drew mostly accurate maps of what they physically experienced.

The averaging subgroup is perhaps the most complex and could have employed a few different strategies in remembering to their staring location, such as an optimal or Bayesian cue combination approach in which they essentially combine or average the self-motion and visual landmark cue (Newman et al., 2023; Vishwanath et al., 2026). It could also be that this averaging subgroup alternated or switched between relying on the illusory intersection and self-motion cues depending on uncertainty, prior knowledge or learning factors (Chen et al., 2017; Harootonian et al., 2022; Qi & Mou, 2024). We did find some evidence for switching, at least in young adults, where we observed a decrease in pointing error across trials in the averaging subgroup (Fig. 5).

For example, a participant could have started the experiment relying on the illusory intersection, but over trials they may have shifted to greater reliance on self-motion cues. A crucial future direction from this task is to determine the nature of cue integration strategies employed when resolving conflicting self-motion and visual intersections cues in a computational model to test some of the above-mentioned theories.

With regard to age, overall, older adults performed statistically indistinguishably from young adults for the accuracy in pointing to the start location and exhibited similar pointing patterns in the illusory condition. The pointing and distance errors in older adults was indistinguishable from young adults (Fig. 2C-D). Both young and older adults showed comparable numbers of participants who showed the cross illusion. The learning characteristics, which were determined by the slope of pointing error across trials, also showed no statistical difference between age groups in both control and illusory conditions (Fig. 4 and 5). Both young and older adults showed a positive correlation between map-drawing accuracy and pointing error (Fig. 6B). Thus, older adults performed in many ways like young adults.

We did find some age effects, however. One consistent age difference was in the variability of pointing error. For the false-intersection pointing error, older adults showed lower trial-to-trial pointing variability compared to the control condition. Additional analyses suggested that older adults in the averaging subgroup of false-intersection condition showed less variable pointing error compared to young adults. Further analysis revealed that older adults did not show improvements in pointing error over trials in the any of subgroups while young adults did. Taken together, the reduced variability in the illusory intersection condition and no discernable improvement across trials, especially in the averaging subgroup, could be consistent with older adults utilizing and combining both the visual and self-motion information and potentially more rigid strategy use than young adults. In other words, older adults may have stuck with their initial weighting or reliance on the visual cues and were less flexible or did not update these estimates across trials when compared to young adults (Kimura et al., 2019; Merhav, 2025). This continued reliance on uncertain or unreliable visual cues has also been suggested to stem from changes in Gamma-aminobutyric acid (GABA) levels and receptor expression and overall decrease in inhibitory regulation among older adults (Merhav, 2025; Rozycka & Liguz-Lecznar, 2017; Zuppichini et al., 2024).

We also found that older adults exhibited greater variability in pointing error in the control or stable condition compared to young adults. One interpretation for the greater variability in pointing patterns in the control condition is noisier path integration estimates in older adults (Harris & Wolbers, 2012; Stangl et al., 2020). However, given that older adults showed greater variability in the stable condition and lower variability in the illusory condition, this finding would also be consistent with sub-optimal weighting of visual landmark cues. This result is supported by previous work suggesting that older adults employ different cue integration strategies compared to young adults (Merhav, 2025; Shayman et al., 2024). For example, a study by Shayman et al., (2024) found that older adults utilized more reliable cues less than young adults, indicating a sub-optimal weighting of cues, which could explain the higher variability in the stable true-intersection condition.

While older adults performed much like young adults in terms of the correlation between pointing error and drawing accuracy, older adults did draw worse maps of the control paths. At the same time, older and young adults showed pointed with comparable accuracy in the true-intersection condition and the accuracy of their pointing correlated with the accuracy of their drawn map. Together, our results do not appear consistent with hypotheses related to a deficit in allocentric navigation. An allocentric, or possibly egocentric deficit, would instead have been reflected in either worse pointing to the start location and worse map drawing or a lower correlation between the two for older adults. Also, if older adults had an overall allocentric deficit, we would have expected fewer accurate maps in the false-intersection condition, particularly for the “accurate” older subgroup, which we did not find. It is important to note, however, that the map-drawings were obtained in virtual reality using a virtual whiteboard and virtual marker, which could have presented challenges in motor control and adaptability differences in older adults compared to young adults.

## Notes

### Competing Interest Statement

The authors have declared no competing interest.

### Funding

- National Institute on Aging: R21 AG081558-01, R01 AG003376
- National Institutes of Health: T32 AG044402

