## Supplementary figure 1 for "Illusory path configurations reveal age-related differences in egocentric pointing variability"

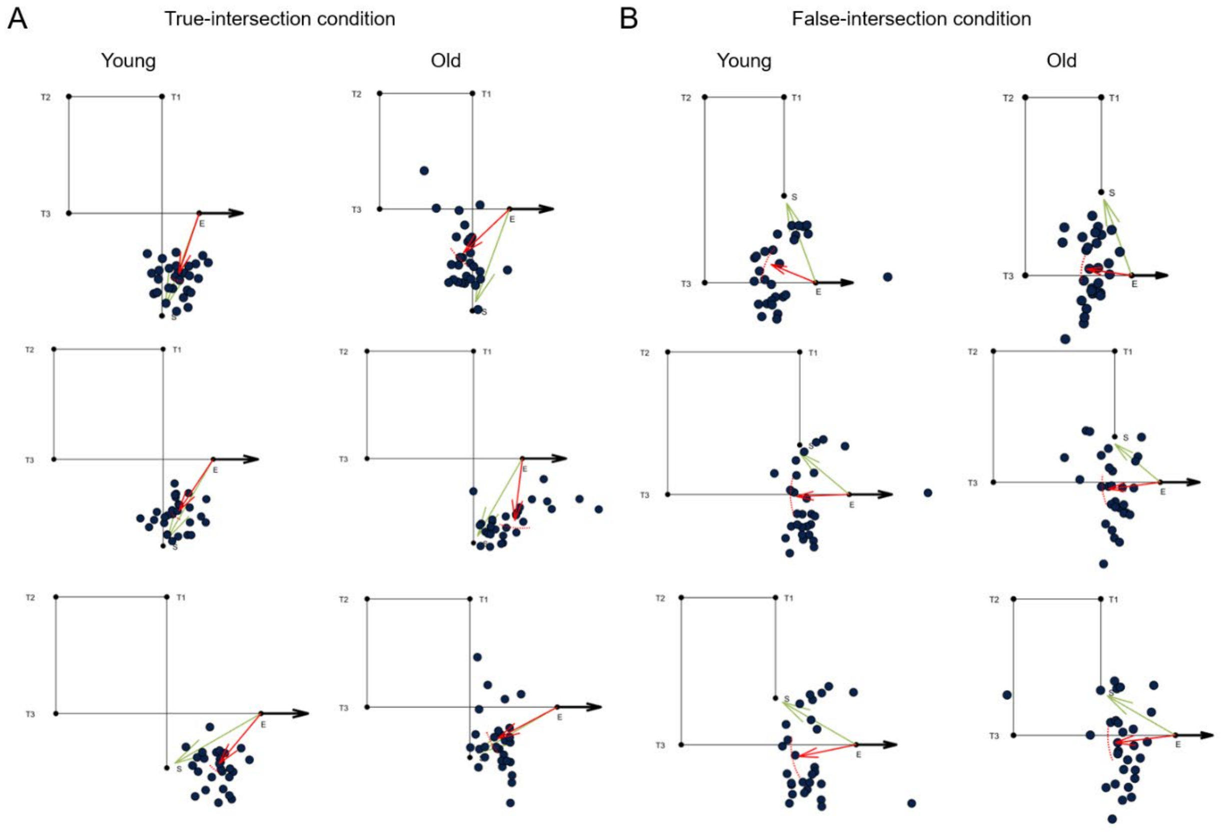


**Supplementary figure 1:** Pointing response in the two Hallway conditions and all unique paths. **A)** Participant's response location (blue dots) on the true-intersection condition averaged across trials in three path configurations for young (left column, n = 28) and older adults (right column, n = 29). The 'S' and 'E' represent the start and end points for the path. The 'T1', 'T2', and 'T3' represent the three turns for the given hallway path. **B)** Same as in A), but for three path configurations in the false-intersection condition.
